# Salt stress disrupts local auxin and COP1 gradients in Arabidopsis apical hooks

**DOI:** 10.64898/2025.12.03.691840

**Authors:** Elizabeth van Veen, Jesse J. Küpers, Xizheng Chen, Yu Him Tang, Thijs de Zeeuw, Kilian Duijts, Scott Hayes, Christa Testerink, Charlotte M. M. Gommers

**Affiliations:** Laboratory of Plant Physiology, Wageningen University & Research, Wageningen, The Netherlands

**Keywords:** skotomorphogenesis, apical hook, salt stress, auxin, CONSTITUTIVE PHOTOMORPHOGENIC1

## Abstract

Seedling establishment is highly sensitive to environmental cues, with light serving as a principal regulator. In darkness, seedlings enter skotomorphogenesis, marked by hypocotyl elongation and apical hook formation, which helps seedlings emerge from the soil to reach the light. Here, we show that salinity impairs soil emergence of *Arabidopsis thaliana* seedlings by inducing a partially photomorphogenic like phenotype in darkness, characterized by reduced apical hook curvature. Within the hook, salt stress diminished differential epidermal cell elongation required to drive hook bending, and reduced both the auxin signalling maximum and PIN3 abundance on the concave side of the hook. Transcriptome analysis revealed that salt and osmotic stresses directly alter organ-specific transcriptional profiles, including the NaCl-mediated repression of *B-BOX DOMAIN PROTEIN 28* (*BBX28*) in hooks and cotyledons. Under control conditions, BBX28 protein accumulated asymmetrically across the hook, but this pattern was lost under salinity. Notably, the ubiquitin ligase CONSTITUTIVE PHOTOMORPHOGENIC1 (COP1) exhibited an opposite gradient to BBX28, which was similarly disrupted by salt stress or by the inhibition of polar auxin transport, presenting auxin as an upstream regulator of COP1 spatial distribution. Together, these findings establish COP1 asymmetry as a novel feature of the apical hook, indicating that spatial regulation—not just absolute levels—shapes COP1 function, and reveal how salinity disrupts hormonal, transcriptional and protein networks to compromise seedling establishment under stress.

**Significance Statement:** Seedling establishment is a critical developmental transition that contributes to survival in the local environment. While light is the primary cue driving de-etiolation, how abiotic stress shapes early development in darkness remains unclear. We show that salinity induces a photomorphogenic-like response in dark-grown *Arabidopsis thaliana* seedlings, disrupting apical hook formation and soil emergence. This response involves disrupted asymmetrical epidermal cell elongation and auxin signalling, repression of *B-BOX DOMAIN PROTEIN 28* (*BBX28*), and loss of a previously unrecognized spatial gradient of the key light regulator COP1 across the apical hook. Our discovery that COP1 is spatially regulated and disrupted by stress, reveals a new dimension of light signalling and developmental control, highlighting how environmental stress constrains seedling establishment.

## Introduction

Plant development is dynamic and responds to local environmental conditions. For newly germinated seedlings, the absence or presence of light results in distinct developmental programs known respectively as skotomorphogenesis and photomorphogenesis (1). For seeds that have germinated underground, skotomorphogenic (or etiolated) growth supports seedlings to outgrow the soil environment. This program is characterized by rapid elongation of the hypocotyl with an apical hook, and tight folding of the cotyledons to protect the apical meristem (2). Contrastingly, upon light exposure hypocotyl elongation is inhibited, the apical hook uncurls, cotyledons open and expand, and root growth rate increases (1, 3). The transition from skotomorphogenesis to photomorphogenesis, known as de-etiolation, is a crucial juncture for seedling establishment. Light stimulated photoreceptor activation initiates rapid changes in molecular and hormonal signalling components, driving this developmental transition.

The identification of mutants that display defective skotomorphogenesis has revealed core repressors of light signalling in seedlings. CONSTITUTIVE PHOTOMORPHOGENIC / DE-ETIOLATED / FUSCA (COP/DET/FUS) proteins selectively degrade photomorphogenesis promoting factors (4). The E3 ubiquitin ligase CONSTITUTIVE PHOTOMORPHOGENIC1 (COP1), often considered the master regulator of light signalling, was among the first photomorphogenesis repressors identified in darkness due to the photomorphogenic phenotype of dark grown *cop1* mutants (5). COP1 activity depends on complex formation with SUPPRESSOR OF PHYA-105 (SPA) proteins in the nucleus (6). Together they target photomorphogenesis promoting transcription factors including ELONGATED HYPOCOTYL5 (HY5), HY5 HOMOGOLOG (HYH), LONG HYPOCOTYL IN FAR-RED 1 (HFR1) and LONG AFTER FAR-RED LIGHT 1 (LAF1) for proteasomal degradation by polyubiquitination in darkness (6-10). The synergistic repression of light signalling pathways by protein complexes and transcription factors is a balancing act, maximizing the seedling’s chances of reaching light before exhausting seed reserves.

Despite the fundamental importance of etiolated growth for seedling survival, knowledge on how abiotic stresses affect this developmental program are limited. Of the stressors threatening agricultural production, soil salinity has emerged as a major issue, affecting 1 381 million ha (Mha), or 10.7% of the total global land area in 2023 (11). Salinity limits plant productivity by imposing osmotic and ionic stress components on plants. As a result, plants experience reduced growth, nutrient imbalances, altered metabolism, and developmental changes or shifts (12-15). To circumvent the detrimental effects of salinity, plants have evolved protective mechanisms including the SALT OVERLY SENSITIVE (SOS) pathway that selectively removes excess intracellular Na^+^ ions (16). Recent studies have revealed crosstalk between SOS and light signalling components. Light activated phytochrome A (phyA) and phyB photoreceptors enhance the kinase activity of SOS2, which in turn promotes phosphorylation-dependent degradation of PHYTOCHROME INTERACTING FACTOR 1 (PIF1) and PIF3 (17). In addition, phyB stability and nuclear accumulation are regulated through phosphorylation by the receptor-like kinase FERONIA (FER), whose activity is inhibited by salinity (18). Recent work has shown that salinity limits hypocotyl elongation at least partially via increased phyB stability (19).

Here, we identify these phenotypic changes as part of a broader suite of traits associated with photomorphogenesis. These changes impair seedling soil emergence, demonstrating the detrimental effect of salinity on establishment in the local environment. We show that under standard conditions, COP1 is asymmetrically distributed within the apical hook, with nuclear accumulation enriched on the concave side. Notably, salt stress abolishes this COP1 gradient and diminishes the auxin signalling maximum on the concave side. Contrary to the constitutively photomorphogenic phenotype of *cop1* mutants, we observe increased total COP1 protein levels under salt stress. Together our findings reveal a direct link between environmental stress, hormonal regulation, and light signalling during early seedling development.

## Results

### Salinity induces a photomorphogenic like phenotype in etiolated seedlings

To assess how salt stress affects skotomorphogenesis, we analysed dark-grown *Arabidopsis thaliana* (Col-0) seedlings exposed to ionic (NaCl, KCl, NaNO_3_, KNO_3_, MgCl_2_) treatments and an osmotic control (sorbitol). In three-day old seedlings we observed distinct phenotypes induced by ionic or osmotic stress (Fig. 1 A-C, *S1 Appendix*, Fig. S1 A). NaCl, KCl, and sorbitol reduced hypocotyl length, with sorbitol causing the strongest inhibition (Fig. 1 B). Hook curvature, however, was specifically reduced by NaCl, KCl, NaNO_3_, and KNO_3_, but not by sorbitol or MgCl_2_, suggesting regulation by monovalent cations (Fig. 1 C; *S1 Appendix*, Fig. S1 A). Since apical hook curvature depends on both formation and maintenance of hypocotyl bending (20), we conducted time-lapse imaging of seedlings grown under control conditions, 75 mM NaCl, 75 mM KCl, and 150 mM sorbitol treatments using infrared imaging. Salinity impaired hook formation, whereas sorbitol extended the maintenance phase (Fig. 1D). Because reduced hypocotyl elongation and apical hook bending are hallmark traits of photomorphogenesis (1, 21), we questioned whether salinity induces additional light associated responses. Indeed, NaCl enhanced root elongation and cotyledon expansion in the dark (*S1 Appendix*, Fig. S1 B and C). We further observed impaired seedling soil emergence (Fig. 1E), mimicking the response of hookless or constitutively photomorphogenic mutants (22, 23).

**Figure 1.**
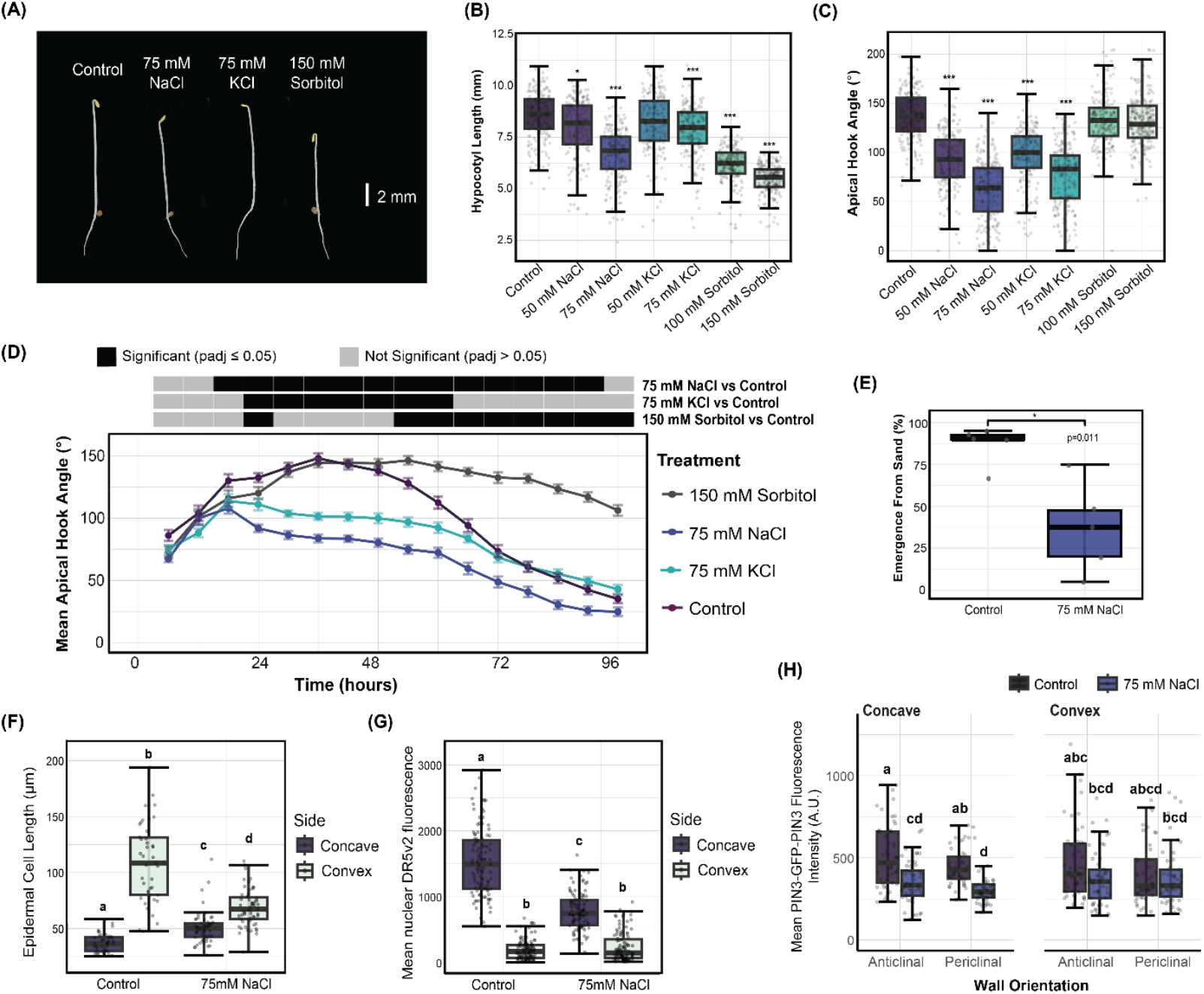
Salinity induces a mild photomorphogenic phenotype in etiolated seedlings, and disrupts the auxin signalling gradient of the apical hook. (A) Representative three-day old dark grown seedlings grown in control medium, or treated with NaCl, KCl or sorbitol. (B and C) Boxplots representing apical hook angles (B) and hypocotyl lengths (C) of three-day old dark grown seedlings treated with different concentrations of NaCl, KCl, and sorbitol. (D) Time-lapse data for apical hook angle following transfer, where points represent mean angle, error bars represent ± SE, n ≥ 40. (E) Percentage of emerging four day old dark grown seedlings from 3 mm sand layer, added to plates post transfer. (F) Boxplot representing epidermal cell size on concave and convex side of apical hook in three day old dark grown seedlings, for 4-5 epidermal cells on concave and convex side, n ≥ 9. (G) Boxplot representing mean nuclear DR5v2:mTurquoise2 signal on concave vs convex side of apical hook in three day old dark grown seedlings, for 12 cells (epidermal and cortex) on concave and convex side, n=9. (H) Boxplot representing plasma membrane localized PIN3-GFP-PIN3 signal intensity on epidermis facing anticlinal and periclinal sides of cortex cell files, on concave and convex sides of the apical hook in two day old seedlings. Boxplots (B, C & E - H) represent the median ± 25% and 50%. Asterisks for panels (B, C and E) refer to significant differences compared to control (* P ≤ 0.05, ** P ≤ 0.05, *** P ≤ 0.001) calculated via Kruskal Wallis followed by posthoc Dunn’s test (B and C), or using independent samples t-test for unequal variances (E). Significance letters (F – H) refer to significant differences (P ≤ 0.05) calculated via Kruskal Wallis followed by posthoc Dunn’s test. Significance for time-lapse data (D) calculated using pairwise t-test with Benjamini-Hochberg (BH) correction for multiple comparisons. All panels refer to seedling assays where seeds germinated on 0.5 MS plates, by exposure to ∼125 μmol/m2/s white light for one hour followed by a further 23 hours in dark before being transferred to respective treatment media.

Because recent research revealed phytochromes as regulators of seedling responses to salt (19), we generated *phyb* and *phyA Arabidopsis* CRISPR knock-out lines (*S1 Appendix*, Fig. S2 A and B). Even though these are severely affected in their response to red and far-red light, respectively (*S1 Appendix*, Fig. S2 C and D), they showed variable skotomorphogenic phenotypes under salt stress and control conditions (*S1 Appendix*, Fig. S2 E and F). All *phyB* mutant lines had significantly shorter hypocotyls under salt stress and control conditions, therefore the reduced sensitivity to salt-inhibited hypocotyl growth observed by Qi et al. (19) was not observed under our experimental conditions. Although *phyB-9* had a higher apical hook angle under salt stress, this line also had the shortest hypocotyl lengths in dark conditions (*S1 Appendix*, Fig. S2F), suggesting delayed development, possibly due to slower germination or the effect of other mutations arising from ethyl methanesulfonate (EMS) mutagenesis. The apical hook angle and hypocotyl lengths of *phyA* mutants behaved the same as Col-0 under salt stress. Together, we show that salt stress promotes photomorphogenic traits in darkness independently of phyA and phyB, compromising seedling emergence and establishment.

### The spatial auxin gradient in the apical hook is reduced by salt stress

Apical hook formation results from differential growth between the concave and convex sides of the hypocotyl, characterized by differences in epidermal cell length between the two sides (24). Based on this, we measured epidermal cell length in the apical hooks of seedlings grown under NaCl to determine whether salinity restricts elongation on the convex side or promotes elongation on the concave side. We observe that salt-treated seedlings exhibited both longer epidermal cells on the concave side, and shorter cells on the convex side compared to non-treated seedlings (Fig. 1F). Considering the typical negative correlation between auxin signalling and epidermal cell elongation within the apical hook (24), we examined how salinity impacts auxin signalling in etiolated seedlings. To investigate spatial changes in auxin signalling across the apical hook under NaCl stress, we used the *DR5v2::mTurquoise2* reporter as a proxy for auxin response. In control seedlings, strong *DR5v2::mTurquoise2* fluorescence was detected predominantly on the concave side of the apical hook (Fig. 1G, *S1 Appendix* Fig. S3A), consistent with the known auxin maximum in this region (20, 25). However, under salinity stress, signal intensity was markedly reduced on the concave side, while fluorescence on the convex side remained largely unchanged (Fig. 1G, *S1 Appendix* Fig. S3A). While increased cell length on the concave side is consistent with reduced auxin signalling under salinity, the difference in cell length on the convex side could not be attributed to altered auxin signalling and may instead reflect auxin-independent effects of salinity on cell expansion. Together these findings show that salinity disrupts differential cell elongation and local auxin signalling in the apical hook.

Asymmetrical auxin transport in the hook relies on the localization of PIN-FORMED auxin efflux carriers (20). Considering that NaCl significantly reduces maximum hook curvature during the formation phase (Fig. 1D), a phenotype that is also exhibited in *pin* mutants (20), we tested for additive effects in *pin3,4,7* seedlings. Salt does not significantly impact hook formation in *pin3,4,7*, and even marginally increased maintenance at 24 hours post germination (*S1 Appendix* Fig. S3B). Although this suggests a role for PINs, *pin3,4,7* seedlings still show reduced hook formation in the first 18 h, which is before the impact of NaCl under our conditions and therefore prevents firm conclusions from timelapse data. To clarify matters further, we assessed the abundance of PIN3-GFP-PIN3 along periclinal and anticlinal cortex cell walls of both hook sides at 24 h, when NaCl has limiting effects on apical hook formation (Fig. 1D). Fluorescence was significantly reduced under NaCl, on the concave but not convex side of the hook (Fig. 1H; S1 Appendix Fig. S3C). Together, these data support the notion that NaCl impacts auxin efflux by reducing the abundance of PIN3 on the concave side of the apical hook under salt stress.

### Salinity Triggers Distinct Transcriptional Responses Across Apical Hooks, Cotyledons and Hypocotyls

Our data show that salt and sorbitol caused overlapping (hypocotyl growth arrest), as well as distinct (hook angle) phenotypes. These differences prompted us to determine whether salinity elicits a general transcriptional response throughout the seedling or distinct responses in different organs/tissues. To that end, we carried out RNA-sequencing of three day old seedlings transferred to 75 mM NaCl, 150 mM sorbitol, or control treatments, dissected into cotyledons, apical hooks, and hypocotyls, (*S1 Appendix* Fig. S4A). Principal component analysis (PCA) revealed that the tissue type had a stronger influence on global gene expression patterns than treatment (*S1 Appendix*, Fig. S5 A). Upon sub-setting of samples into their respective dissected seedling parts however, a clear stress response was observed (*S1 Appendix*, Fig. S5 B-D). Differential gene expression analysis further revealed that the response to salinity or osmotic stress differed across seedling parts, with the highest number of differentially expressed genes (DEGs) in hypocotyls (Fig. 2A, *Appendix* Table S1). Few DEGs were identified for the cotyledons of sorbitol treated seedlings, suggesting the osmotic stress effect on gene expression increases in more basal tissues (Fig. 2A).

**Figure 2.**
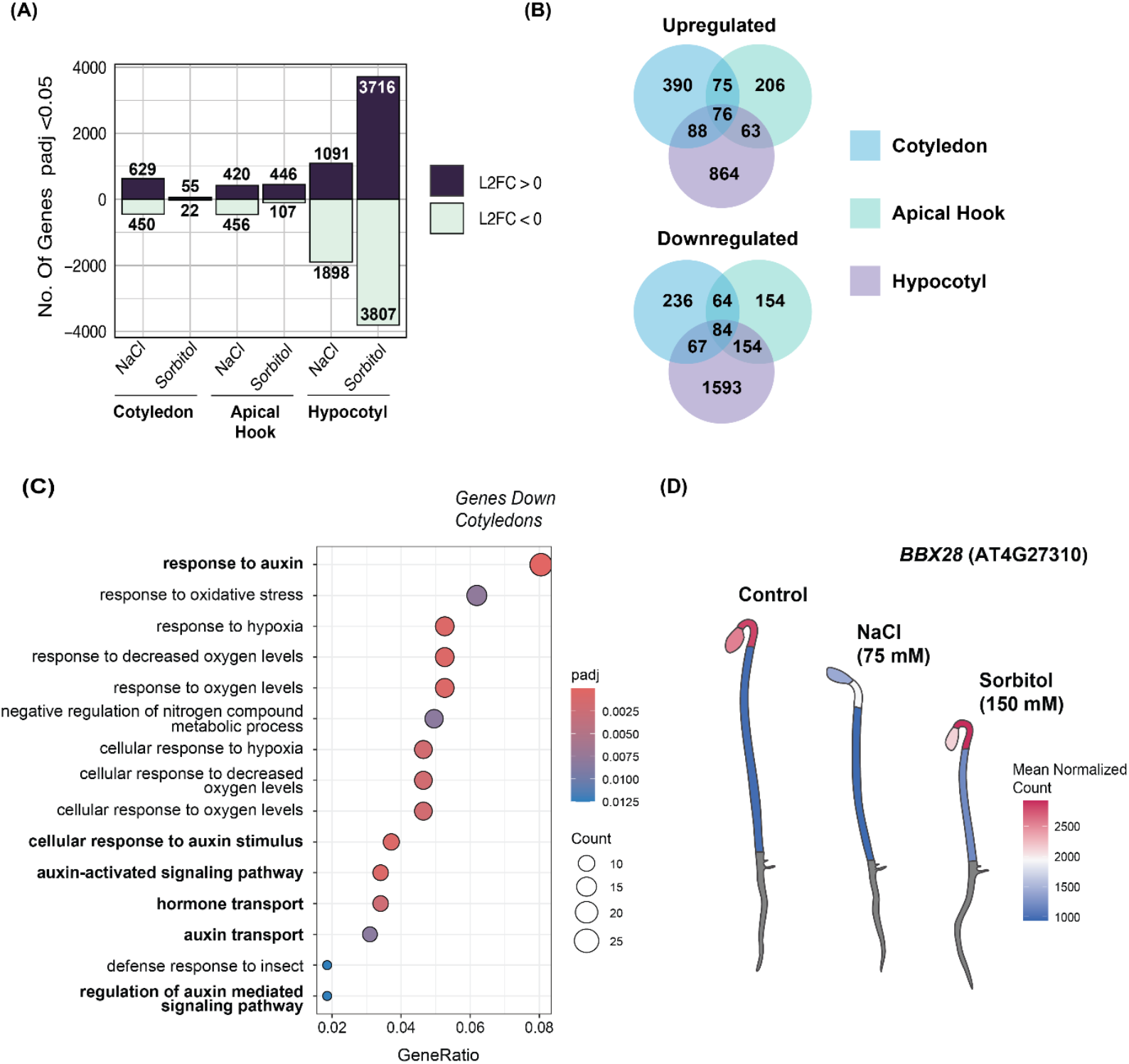
salinity induces distinct transcriptional response in the cotyledons, apical hooks, and hypocotyls of etiolated seedlings. (A) Total number of genes differentially up- or downregulated in response to 75 mM NaCl or 150 mM sorbitol in different seedling organs (padj <0.05, Log2FC ≠ 0). (B) Venn diagram depicting overlap in differentially expressed genes in response to 75 mM NaCl, across cotyledons, hypocotyls and apical hooks (padj <0.05, Log2FC ≠ 0). (C) Bubble plot representing GO enrichment analysis (all domains) for RNA-seq identified DEGs downregulated in cotyledons in response to 75 mM NaCl in three day old etiolated seedlings. Top 15 enrichment terms shown. (D) GGPlantmap expression map showing relative transcript abundance (DESeq2 normalized count) of BBX28.

When assessing overlap in salt-responsive DEGs, only 3.5% of upregulated and 3% of down-regulated genes were shared across the organs (Fig. 2B). The low overlap between the apical hook and the remainder of the hypocotyl (particularly for upregulated DEGs) was striking given that these regions belong to the same organ. These observations suggest that salinity activates distinct gene expression programs in different parts of the seedling. Next, we performed Gene Ontology (GO) analysis to find patterns among differentially expressed genes by salt and sorbitol in the different tissues (*Appendix* Table S2, Table S3). Interestingly, among salt-repressed genes in the cotyledons we found a strong over-representation of auxin-responsive genes (Fig. 2C). Given that auxin transport from the cotyledons and meristem to the hypocotyl is required for hook formation (26), reduced auxin signalling in the cotyledons likely contributes to the diminished auxin response aximum in the hook (Fig. 1G), indicating that salinity disrupts auxin flux and coordination between these tissues.

To exemplify spatial differences in the expression of specific genes across etiolated seedling parts, we generated custom electronic Fluorescent Pictographs (eFP)-style expression maps (using GGPlantmap (27)). Notably, we observed salt-specific downregulation of *SUCROSE-PROTON SYMPORTER 1* (*SUC1*) in cotyledons (*S1 Appendix* Fig. S4B) and *B-BOX DOMAIN PROTEIN 28* (*BBX28*) in cotyledons and apical hooks (Fig. 2D). Contrastingly, *LONG HYPOCOTYL IN FAR-RED 1* (*HFR1*) was specifically upregulated by salt in hypocotyls (*S1 Appendix* Fig. S4C). Hypocotyl tissue is typically under-represented in total RNA extracts. Our data support a reconsideration of bulk seedling RNA sampling strategies when targeting transcripts not abundantly expressed in the cotyledons.

### NaCl represses *BBX28* expression, and disrupts its protein gradient in the apical hook

Transcriptional analyses revealed a reduction in *BBX28* transcripts in response to salinity in the cotyledons and apical hooks of etiolated seedlings (Fig. 2D, Fig. 3A). Given that BBX28 is a known inducer of hook curvature in *Arabidopsis*, and dark-grown *bbx28* mutants have more open apical hooks than wild-type (28), we questioned whether over-expressing *BBX28* could restore apical hook curvature under salt stress conditions. To assess this, we screened the apical hook phenotype of previously described *BBX28* over-expressing line (*35S::YFP-BBX28 #18*) and mutant (*bbx28-3*) (28). Over-expression of *BBX28* increased hook angle under salt stress as well as control conditions, whereas the *bbx28-3* mutant exhibited reduced response to salinity compared to Col-0 (Fig. 3B). These data suggest that *BBX28* comprises one component of a salt-responsive transcriptional network that induces mild photomorphogenesis in dark grown seedlings.

**Figure 3.**
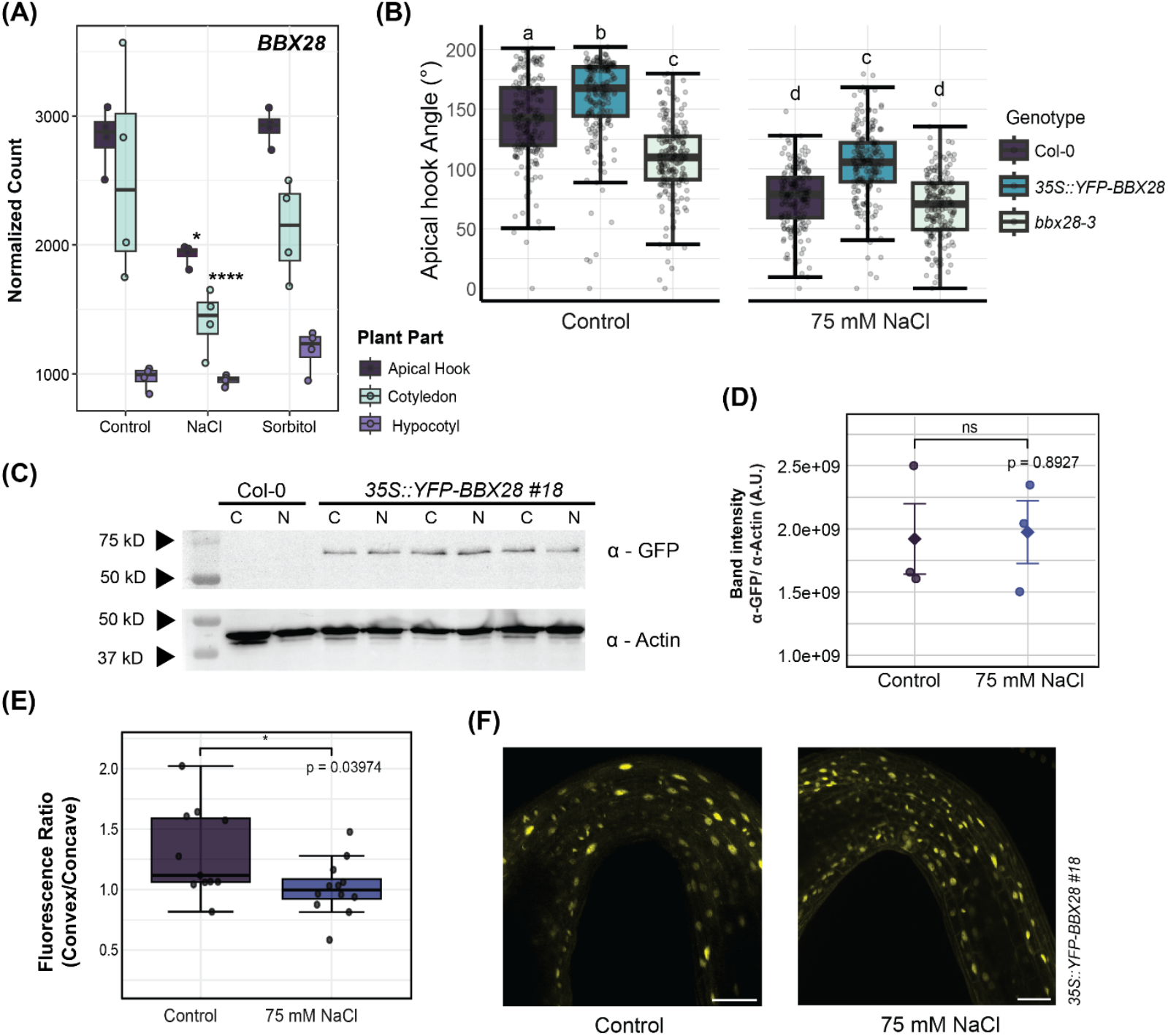
Salinity alters expression and spatial accumulation of BBX28 in apical hooks. (A) Absolute expression of BBX28 from RNA-sequencing of three-day old etiolated seedlings in normalized reads calculated using DESeq2. (B) Boxplot representing apical hook angles of three-day old dark grown seedlings treated with 75 mM NaCl or grown on control media. (C) 35S::YFP-BBX28 protein levels analysed in protein extracts from 40 three-day old etiolated seedlings. Col-0 used as negative control. YFP-BBX28 protein was detected using α-GFP antibody and α-actin was used as a reference. C = control, N = 75 mM NaCl. (D) Boxplots show the YFP-BBX28/actin ratio of band intensity in (C). (E) Boxplot representing ratio in mean nuclear YFP-BBX28 signal intensity on convex / concave side of the apical hook. Mean nuclear signal was calculated for 4-8 nuclei on concave and convex sides of each hook, n=11 (Control) n=12 (75 mM NaCl). (F) Representative confocal images of YFP-BBX28 in apical hooks of three day old etiolated seedlings treated with 75 mM NaCl or grown on control media. Scale bars = 50 μm. Boxplots (A, B, and D) represent the median ± 25% and 50%. Significance calculated using DESeq2 pairwise comparison (A), Kruskal-Wallis and post-hoc Dunn’s test (B), and independent samples t-test (D). Asterisks indicate significant differences compared to control (* P ≤ 0.05, ** P ≤ 0.05, *** P ≤ 0.001) (A, D, E) All panels refer to phenotype data where seeds germinated on 0.5 MS plates, by exposure to ∼125 μmol/m2/s white light for one hour followed by a further 23 hours in dark before being transferred to respective treatment media.

To assess BBX28 protein stability under salt, we carried out immunoblotting of three day old *35S::YFP-BBX28* #18 seedlings grown under control conditions or 75 mM NaCl. YFP-BBX28 levels remained consistent across treatments (Fig. 3C and D). To address spatially restricted stress responses, we investigated the abundance of YFP-BBX28 in the apical hooks of three-day old etiolated seedlings using confocal microscopy. Under control conditions, we newly identified asymmetric nuclear accumulation in YFP-BBX28 with higher signal intensity on the convex side of the apical hook. Contrastingly, in NaCl-treated seedlings nuclear fluorescence was uniformly abundant across the hook (Fig. 3E and F). These data suggest that under control conditions, BBX28 undergoes localized degradation or nuclear exclusion on the concave side of the apical hook, which is lost under salt stress.

### Salinity and NPA similarly disrupt asymmetric nuclear COP1 accumulation across the apical hook

BBX28 undergoes COP1 dependent 26S proteasomal degradation in darkness (28). Given that BBX28 protein accumulates asymmetrically in the apical hook, we examined whether COP1 exhibits asymmetric distribution as well. Confocal imaging of native promoter driven *mCherry-COP1* in 3-day-old etiolated seedlings revealed an inverse gradient, with COP1 enriched on the concave side (Fig. 4A and B, 3E and F). This local accumulation may promote proteasomal degradation of targets such as BBX28. Notably, salinity abolished the COP1 gradient, paralleling the loss of BBX28 asymmetry (Fig. 3E and F, 4A and B). This finding suggests that a non-asymmetric distribution of BBX28 across the hook under salt stress may be a consequence of the loss of COP1 asymmetry. Considering that increased and more uniform COP1 accumulation was present in salt treated apical hooks, we assessed the sensitivity of the *cop1-4* mutant to salinity during the formation phase (24hr post transfer). *cop1-4* seedlings had significantly less curved hooks when compared to Col-0 under control conditions, but not under salt (Fig. 4C). These data reflect a requirement for functional COP1 to form an apical hook under control conditions. In the absence of this however, salt stress has no additional impact on the apical hook angle, consistent with a requirement for COP1 in salt responsive phenotypic change. mCherry-COP1 levels from whole seedling protein extracts were slightly higher under salt stress than control conditions, whereas *COP1* gene expression is unaffected across all tissues (Fig. 4D-F). Together, these data show that COP1 is post-transcriptionally upregulated under salt stress, and disrupting the COP1 gradient either through salt treatment (Fig. 4A and B), or impairing COP1 function via mutagenesis (Fig. 4C) are associated with reduced apical hook curvature.

**Figure 4.**
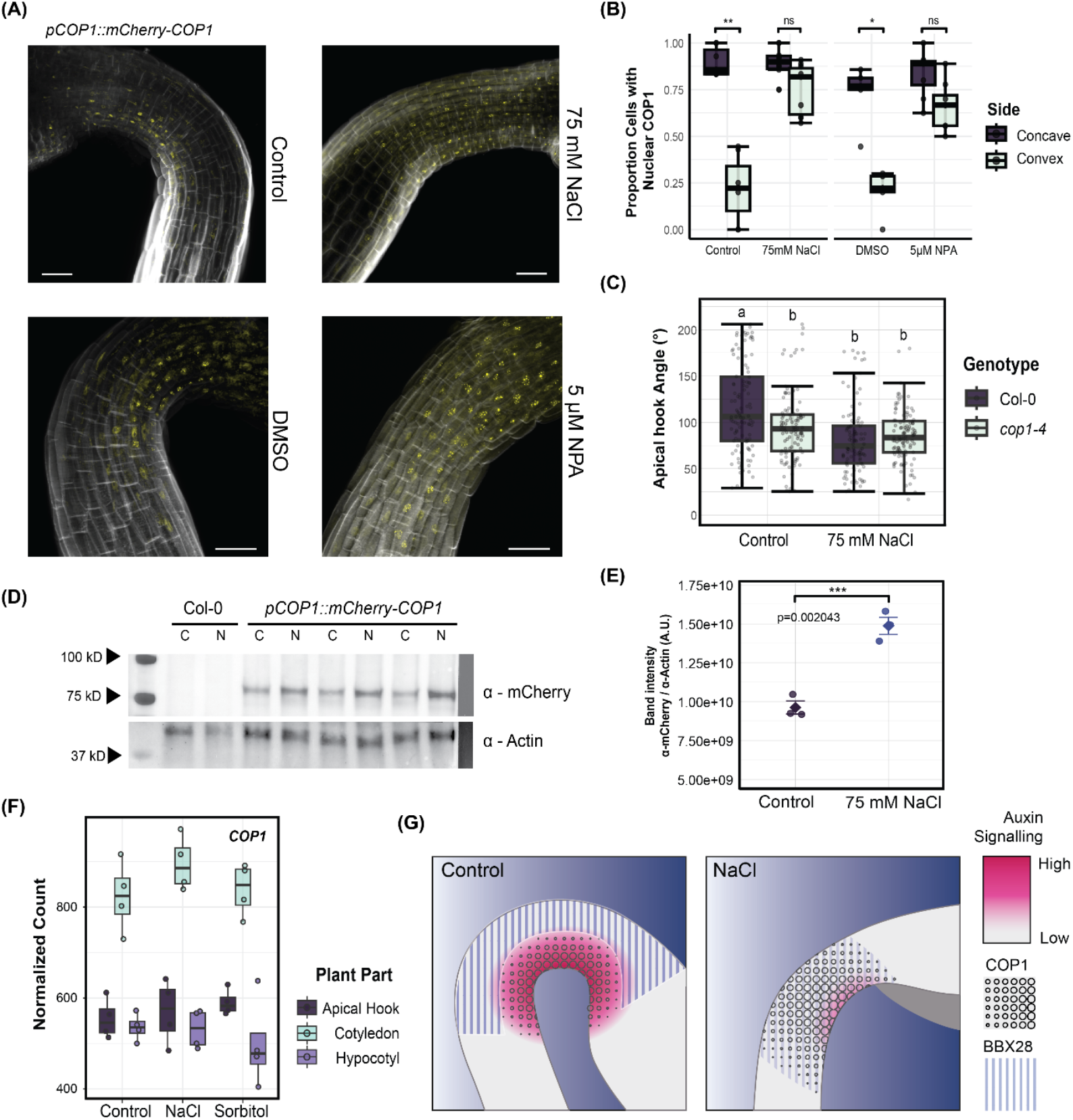
NaCl and NPA similarly increase homogeneity in mCherry-COP1 nuclear accumulation across the apical hook of etiolated seedlings. (A) Confocal images of mCherry-COP1 (yellow) in apical hooks of three day old fixated etiolated seedlings grown under control conditions, mock (DMSO), 75 mM NaCl, or 5 μM NPA. Cell walls are stained with calcofluor white (white). Scale bars = 50 μm (B) Proportion of cells in file of cortex cells with nuclear mCherry-COP1 signal divided by total number of visible cells for concave and convex sides of apical hook. (C) Boxplot representing apical hook angles of two-day old dark grown seedlings treated with 75 mM NaCl or grown on control media. (D) mCherry-COP1 protein levels analysed in protein extracts from ∼40 three-day old etiolated seedlings. Col-0 used as negative control. mCherry-COP1 protein was detected using α-mCherry antibody, α-actin was used as a reference. C = control, N = 75mM NaCl. (E) Quantified mCherry-COP1 levels in terms of band intensity divided by band intensity of Actin. Significance calculated using t-test with homogenous variances. (F) Absolute expression of COP1 from RNA-sequencing of three-day old etiolated seedlings in normalized reads calculated using DESeq2. (G) Summarizing schematic depicting how NaCl impacts gradients in nuclear COP1 and BBX28 accumulation, and auxin signalling gradients within the apical hook of etiolated seedlings. Boxplots (B, C, E & F) represent the median ± 25% and 50%. Significance calculated using Kruskal-Wallis and post-hoc Dunn’s test whereby asterisks indicate significant differences compared between concave and convex sides (* P ≤ 0.05, ** P ≤ 0.05, *** P ≤ 0.001) (B), aligned rank transform ANOVA with Holm-corrected post-hoc comparisons (sig. P ≤ 0.05) (C) independent samples t-test (E), and DESeq2 pairwise comparison (F). All panels refer to phenotype data where seeds germinated on 0.5 MS plates, by exposure to ∼125 μmol/m2/s white light for one hour followed by a further 23 hours in dark before being transferred to respective treatment media for a further 48 hours (A, B, D, E & F), or 24 hours (C).

Interestingly, the COP1 accumulation in the concave side of the hook under control conditions mimics that of the auxin response as observed with the DR5v2 reporter (Fig. 1G, S1 *Appendix* Fig. S3A) (25). Considering that salinity reduces auxin signalling and PIN3 abundance on the concave side of the hook (Fig. 1H; S1 Appendix Fig. S3C) we were prompted to assess the interaction between auxin and COP1 by disrupting polar auxin transport with NPA. Strikingly, NPA treatment reduces the gradient in nuclear COP1 accumulation similar to NaCl (Fig. 4A and B). These findings show that restriction of COP1 accumulation to the concave side of the apical hook requires polar auxin transport, which we have previously shown to be impacted by salt stress.

## Discussion

In *Arabidopsis thaliana*, salt stress delays developmental transitions and suppresses growth, including seed germination (17, 29), primary and lateral root elongation (30, 31), and flowering (32). Here, we observe that salinity induces de-etiolation like physiology in dark grown seedlings, characterized by reduced hypocotyl length and apical hook curvature, but increased root growth and cotyledon area (Fig. 1 A-C, *S1 Appendix* Fig. S1 A-C). Though these changes indicate a promotion of photomorphogenic onset, this is detrimental in the soil environment with salt stressed seedlings exhibiting reduced emergence (Fig. 1E). The reduced hypocotyl elongation and apical hook bending response has been independently confirmed (19). Unlike Qi et al. (19), we did not detect phyB involvement in hypocotyl inhibition as newly generated *phyB* mutants did not have longer hypocotyls when exposed to salt stress under our experimental conditions (*S1 Appendix* Fig. S2 E and F). Collectively, these results indicate that salinity can promote de-etiolation in the dark independently of phyB.

In this study, we identified differences in the transcriptional response to salinity across the cotyledons, apical hooks, and hypocotyls of etiolated seedlings (Fig. 2F and G). A similar dissection approach was previously used to identify *BUB3*.*1* as an organ-specific transcriptional target of PIFs, with expression enriched in the cotyledons and apical hook (33). Notably, this study focussed on PIF-dependent regulation of etiolated development, and identified highest fraction of DEGs in the cotyledons and least in the hypocotyl (33). We now identified the most pronounced transcriptional responses to salt and osmotic stress were observed in the hypocotyl (Fig. 1F), suggesting that stress cues elicit organ specific transcriptional responses that differ from those associated with light-dependent development.

Among stress-responsive DEGs, *BBX28* was specifically repressed by salt—but not osmotic stress—in the apical hook and cotyledons (Fig. 2D, Fig. 3A). Given the open-hook phenotype of *bbx* mutants (30), this repression likely contributes to salt-induced reduction of hook curvature. Consistently, *BBX28* overexpression enhanced curvature, whereas *bbx28-3* mutants resembled wild type under NaCl (Fig. 2B). Because both lines still responded to salt, BBX28 likely acts within a broader transcriptional network that promoties mild photomorphogenesis under salinity. B-box proteins, long known as light-signalling components, are increasingly recognized as regulators of abiotic stress (34-37). For instance, overexpression of *BBX24* (formerly *SALT TOLERANCE, STO*) promotes root growth and germination under saline conditions (38, 39). Recent work has shown that the *BBX28* locus comprises a natural antisense transcript (NAT) pair that contributes to cold acclimation and freezing tolerance (40). Our data support this broader role for B-box proteins as context-specific regulators integrating developmental and environmental responses.

*BBX28* is subject to both transcriptional and post-translational regulation. While total protein levels remained unchanged between control and salt-treated conditions (Fig. 2C and D), we observed a striking asymmetry in nuclear YFP-BBX28 signal intensity across the apical hook under control conditions, with enhanced accumulation on the convex side. This spatial distribution resembles patterns reported for other hook regulatory components. For example, a similar gradient in nuclear accumulation was observed for CELL CYCLE SWITCH PROTEIN 52 A2 (CCS52A2), which promotes endoreplication on the convex side of the hook (41). Likewise, *SMALL AUXIN UPREGULATED RNA 57* (*SAUR57)* expression is enriched on the convex side, thereby spatially restricting the activity of the phosphatase *PP2C-D1* to the concave side (42, 43).

The newly observed confinement of nuclear COP1 to the concave side of the hook under control conditions (Fig. 4A and B), suggests that the asymmetric BBX28 gradient is established through localized degradation via the 26S proteasome. Supporting this, overexpression of *BBX28* rescues the short hypocotyl—but not the open hook—phenotype of *cop1-6* mutants in the dark, and promotes elongation in the light, when COP1 is inactive (9). These findings support a model in which BBX28 promotes cell elongation throughout the hypocotyl, while its exclusion from the concave side of the hook may be necessary to maintain asymmetric growth and curvature under control conditions. Under salt stress, the disruption of both COP1 and BBX28 gradients (Fig. 3E and F, Fig. 4A and B) further supports the idea that COP1 regulates the spatial distribution of *BBX28* in the apical hook.

Our finding that NaCl increases both total COP1 protein levels and its nuclear localization across the apical hook (Fig. 4A and B) appears paradoxical given the constitutively photomorphogenic phenotype of *cop1* lines. These results point to a more nuanced, context-dependent role for COP1 in integrating skotomorphogenic development with environmental stress responses then what was previously assumed. This view is supported by recent research showing that COP1 function is both organ and tissue specific. For example, COP1 targets the transcription factor BES1 for degradation in the cotyledons but stabilizes it in the hypocotyl under shade or warm temperatures (44). Moreover, COP1 was shown to exhibit high tissue specificity in roots, localizing to trichoblasts within the elongation zone, but to the endodermis of the differentiation zone (45). Similar spatial specificity has been observed for other light-regulated proteins, such as the PIFs, which exhibit cell-type-specific transcriptional control—for example, activating auxin-responsive genes in endosperm tissues (46), but inhibiting stress responsive gene expression in guard cells (47).

Past research has implicated COP1 as an upstream regulator of auxin signalling, influencing its biosynthesis, polar transport, and metabolism (48, 49). For example, *cop1* mutants fail to exhibit the typical increase in auxin levels observed under shade conditions in wild-type seedlings (50), and the high-temperature induction of auxin biosynthesis gene *YUCCA8* (*YUC8*) is lost in *cop1-4* (51). In terms of polar auxin transport, the dark induced reduction in *PIN1* expression in hypocotyls is attenuated in *cop1* mutants, whereas PIN1 and PIN2 protein abundance at the plasma membrane are also regulated by COP1 (52). Our observation that salt stress reduces PIN3 abundance on the concave side of apical hook (Fig. 4F) is consistent with these findings and indicates that the salt-induced increase in COP1 level may contribute to PIN3 destabilization or degradation. However, the concurrent loss of the COP1 gradient and increase in the nuclear COP1 levels across the hook by NPA (Fig 3A and B) point to a more complex regulatory network. Auxin-dependent control of COP1 activity is supported by a recent study that found the high auxin levels in hypocotyls under shade conditions, induce nitric oxide levels, which promote COP1 activity through S-nitrosylation (53). Our findings now suggest a feedback mechanism, whereby the chemical disruption of auxin transport triggers COP1 activity, which in turn may modulate auxin transport components.

Taken together, our findings reveal that salinity elicits specific spatial regulatory changes in etiolated *Arabidopsis* seedlings and challenge the prevailing view of salt stress as inhibitory to development. By combining organ-specific transcriptomics with spatial protein localization and DR5v2 imaging, we identify *BBX28* as a salt-repressed hook regulator. Moreover, we show salt disrupts asymmetric gradients in both BBX28 and COP1, as well as auxin signalling across the hook, highlighting a novel point of integration between light responsive and salt stress signalling networks (Fig. 4G). Importantly, we show that the salt-induced loss of hook curvature impairs seedling emergence from the soil, underscoring the ecological and agronomic relevance of apical hook integrity for establishment under saline conditions. Altogether, we reveal the apical hook as a critical site of light, hormone, and stress integration that safeguards early establishment under challenging environments.

## Materials and Methods

### Plant materials and growth conditions

*Arabidopsis thaliana* plants used for experimental work were in the ecotype Columbia-0 (Col-0) wild-type background. The *cop1-4* (54), *phyB-*9 (55), *phyA-211* (56), and *pin3-3 pin4-101 pin7-102)* (57) mutant lines are described previously. Transgenic lines carrying *35S::YFP-BBX28 (#18)* (28), *pCOP1::mCherry-COP1 (45)*, and C3PO (58) constructs were all in Col-0 background as already described, whereas *pPIN3::PIN3-GFP-PIN3* was in *pin3-3* (Col-0) mutant background (20). Seeds were surface-sterilized in 0.05% Triton X-100 for 5 min, then in 1:1 household bleach:0.05% Triton X-100 for 10 min, and rinsed four times with sterile water. Sterilized seeds were sown on 50 µM nylon mesh (Sefar BV) over half-strength MS medium (Duchefa) with 0.8% plant agar (Duchefa), pH 5.7. After 4 d stratification at 4 °C in darkness, seeds were exposed to ∼125 μmol m−^2^ s−^1^ white light at 21 °C for 1 h (1-4 hours for *phy* mutants), then wrapped in foil and grown vertically in darkness at 21 °C for 23 h. Seedlings were subsequently transferred under safe green light to new 0.5× MS plates with or without treatments (see below) and grown in darkness for 48 h unless otherwise stated.

### Skotomorphogenic phenotype analyses

For single timepoint experiments (excluding cotyledon size), photos of seedlings were taken with a Canon EOS R10 camera one to two days after transfer to treatment media (three days old in total). Cotyledons were imaged using a Leica DM2500 microscope. For time-lapse experiments, plates were placed in a previously described infrared imaging set up directly after transferring to treatment media (31, 59). Phenotype analyses were carried out in Fiji (60). Hypocotyl lengths were measured using the segmented line tool, cotyledon size was measured using the polygon toon, and apical hook angle was measured using the angle tool as according to a previously (61).

### Stress and pharmacological treatments

For salt stress experiments, 50 mM or 75 mM of NaCl (Fluka), KCl (Amresco), NaNO_3_ (Sigma), KNO_3_ (Sigma), and MgCl_2_ (Sigma) were added to 0.5 MS media. For osmotic stress experiments, 100 mM or 150 mM of sorbitol was added to 0.5 MS media. Stock solutions of N-1-naphthylphthalamic acid (NPA; Supelco) were made by dissolving directly in dimethyl sulfoxide (DMSO; Sigma) to reach a stock concentration of 10 mM, which was later added to media to achieve a working concentration of 5 µM. Equal volumes of DMSO (0.05%) were added to 0.5 MS as mock treatment.

### Generation of CRISPR *phyA* and *phyB* alleles

The CRISPR alleles of *PHYA* and *PHYB* were generated using CRISPR-zCas9i as described previously (62). Four gRNAs spanning the coding region were designed for both *PHYA* and *PHYB* using CHOP-CHOP (63). In addition, potential off-targets were checked using BLAST (Araport11). The gRNAs were GoldenGate assembled into shuttle vectors pDGE332, pDGE333, pDGE335 and pDGE337 (Table S4). Loaded shuttle vectors were subsequently used to assemble into transformation vector pDGE347. The presence of the correct gRNAs was confirmed by Sanger sequencing of the loaded transformation vector. *Agrobacterium tumefaciens* AGL-0 carrying the loaded transformation vector were used to transform Col-0 with the floral dipping method. T1 transformants were selected based on seed fluorescence (FAST marker) (MZ10-F Microscope, Leica) and transferred to soil. T1 plants were screened for mutations by PCR and Sanger sequencing using genotyping primers that cover all gRNA positions (Table S5). Successful *PHYB* T1 transformants were also identified by choosing plants with elongated hypocotyl and petioles. Only non-fluorescent T2 (absence of Cas9) seeds were utilized for further selection. In T3 generation, absence of Cas9 was confirmed by PCR genotyping using Cas9 specific primers. Homozygous and Cas9-free *phyA* and *phyB* mutants were used for experiments

### Protein isolation and immunoblotting

Three-day-old *pCOP1:mCherry-COP1*, Col-0, *cop1-4, 35S::YFP-BBX28* seedlings were harvested in the dark under a green safe-light. Forty etiolated seedlings per sample were flash-frozen in liquid nitrogen and stored at –80 °C in 2 mL safelock tubes with steel ball bearings until extraction. Samples were ground twice at 25 Hz for 30 s, resuspended in 80 μL protein extraction buffer (for *pCOP1:mCherry-COP1*: 50 mM Tris pH 7.5, 200 mM NaCl, 4 M urea, 0.1% Triton X-100, 5 mM DTT, cOmplete protease inhibitor cocktail tablet (Roche), for *35S::YFP-BBX28*: 0.12 M Tris-HCl pH 7.5, 5 mM EDTA pH 8, 4 % SDS, 5 % glycerol, 50 mM DTT, cOmplete protease inhibitor cocktail tablet (Roche)), and centrifuged at 13,000 rpm at 4 °C for 15 min. Supernatants were mixed with 5× laemmli buffer (300 mM Tris-HCl pH 6.8, 5% SDS, 50% glycerol, 100 mM DTT, 0.05% bromophenol blue), heated to 95 °C for 5 mins and kept on ice. Proteins were separated on 7.5% stain-free polyacrylamide gels (Bio-Rad) for mCherry-COP1, or 12% SDS-PAGE gels for YFP-BBX28, and transferred to nitrocellulose membranes using the Trans-Blot Turbo system (Bio-Rad). Membranes were blocked in 5% milk/TBST for 1 h, then incubated overnight at 4 °C with primary antibodies: anti-mCherry (1:1000 Abcam ab167453), anti-GFP (1:1000, ChromoTek 3H9) or anti-actin (1:1000 Santa Cruz sc-47778). After washing, membranes were incubated with anti-rat (1:5000, Agrisera AS10 1224), anti-rabbit (1:2000 PhytoAB PHY6000) or anti-mouse (1:2000 PhytoAB PHY6006) secondary antibodies for 1 h at room temperature, washed again, and visualized with Clarity ECL substrate on a ChemiDoc system (Bio-Rad). Signal intensity of protein bands was quantified using Image Lab software, volume tool (Bio-Rad, version 6.1).

### Confocal Microscopy and Sample Preparation

Fixation was carried out on three day old dark harvested *pCOP1::mCherry-COP1* seedlings by immersion in 4% paraformaldehyde (in PBS) for two hours under a light vacuum in the dark. Seedlings were subsequently washed with PBS three times, and then immersed in ClearSee solution (64) in the dark for one week. Cell walls were counterstained with 0.05% calcofluor white (Megazyme) in ClearSee for one hour and subsequently destained in ClearSee for one hour. Imaging was carried out by mounting the seedlings on slides with ClearSee. Cell wall staining of live tissues was conducted by immersing Col-0 seedlings in a 20 µg/mL propidium iodide (Sigma) solution for seven minutes, followed by washing with water three times. *C3PO, PIN3-GFP-PIN3*, and *35S::YFP-BBX28*, and PI stained Col-0 imaging was carried out on live tissue of age described in figure legends. Confocal images were acquired using a Stellaris 5 Confocal LSM Leica microscope, with either a 40x oil immersion lens or a 20x dry lens. Microscope setting were as follows: mCherry-COP1: excitation 587 nm, emission ∼592-750 nm, calcofluor white: excitation 350, emission ∼410-480, DR5v2::mTurqoise2: excitation 448, emission ∼459-480, YFP-BBX28: excitation 514, emission ∼525-548, PI: excitation 549, emission ∼591-655 and PIN3-GFP-PIN3: excitation 485, emission ∼490-528.

### Confocal Microscopy Analyses

All confocal microscopy quantification were carried out using Fiji (60). Epidermal cell lengths were measured using the segmented line tool. PIN3-GFP membrane fluorescence was measured by generating max projections with the sum-slice function for Z stacks of the first cortex layer, followed by mean fluorescence intensity along membranes using the segmented line tool, line width = 3. Nuclear fluorescence intensities were measured by generating max projections with the sum-slice function for Z stacks that are approximately one cell layer thick. Nuclei were selected using the selection brush tool. To account for nuclei with low *DR5v2::mTurqoise2* fluorescence, the C3PO reporter was used (65), and nuclei were selected using *mD2-ntdTomato* channel.

### Seedling Dissection and Extraction of RNA from Dissected Seedlings

Three day old etiolated seedlings were grown as described above. Dissections were carried out under dark conditions with a green safe-light. Etiolated seedlings were rinsed in a 1:1 solution of RNA-later (invitrogen) and RNAse free water, before separating cotyledons, apical hooks, and hypocotyls (Fig S2A) with a scalpel. Dissected organs were then placed in pre-cooled 2ml safelock Eppendorf tubes containing two 1/8” steel ball bearings (Weldtite, 3906141), and flash frozen in liquid nitrogen. Each tube contained organs from 30-45 seedlings, and were ground frozen for 30s at 25 Hz twice. RNA-isolation was carried out using the RNEasy Plant Mini kit (Qiagen) with on-column DNAse treatment, in accordance with manufacturer instructions. Three samples were combined during addition of RLT buffer, whereby the buffer was transferred from one tube to the next, until the lysate of 90-125 seedlings was suspended in the buffer to be loaded into the QIAshredder spin column.

### RNA-Sequencing Analysis

RNA poly-A enrichment library preparation and sequencing were carried out by Novogene using the illumina NovaSeq 6000 platform. Read quality checking, trimming, mapping and quantification against the TAIR10 genome (66) were conducted using a NextFlow nf-core/rna-seq pipeline v3.9 (67). Differential gene expression analyses were conducted in R using DESeq2 with a minimum read cutoff of ≥5 reads in ≥9 samples (68), with subsequent LFC shrinkage (69) to correct for variability in lowly expressed genes. GGPlant pictograms were generated using the GGPlantmap package in R (27). Gene Ontology enrichment was performed separately for up- and down-regulated genes in each organ under NaCl treatment using clusterProfiler (70) in R with TAIR annotations, considering all GO domains and controlling false discovery rate at 0.05.

### Statistical analysis and data representation

Statistical analyses were carried out in RStudio (version 4.4.1) Information regarding used statistical methods are provided in the figure legends. For detailed results of the conducted analyses, refer to Supplementary table S6. All graphs were generated using the R package ggplot2. Maximum intensity projections of confocal Z-stacks were made in Fiji.

## Acknowledgements

We are grateful to Sanne Matton and Ronald Pierik for sharing Arabidopsis *pin* mutants and *PIN3-GFP-PIN3* lines, to Dongqing Xu for sharing Arabidopsis *bbx28* mutants and *35S::YFP-BBX28* lines, and to Karen Halliday for sharing *phyB-9* and *phyA-211* lines. We acknowledge support from the Graduate School for Experimental Plant Sciences (EPS), Young Tenure Track grant awarded to C.M.M.G, the Dutch Research Council (NWO), Vici (VI.C.192.033) and ENW-KLEIN (OCENW.KLEIN.421) awarded to C.T., the Wageningen Graduate School Postdoc Talent grant to S.H, and the contribution of the Sector Plan Biology of Wageningen University funded by the Dutch Ministry of Education, Culture, and Science, to C.M.M.G.

## References

1. C. M. M. Gommers, E. Monte, Seedling Establishment: A Dimmer Switch-Regulated Process between Dark and Light Signaling. Plant Physiol 176, 1061–1074 (2018).

2. A. Seluzicki, Y. Burko, J. Chory, Dancing in the dark: darkness as a signal in plants. Plant Cell Environ 40, 2487–2501 (2017).

3. A. A. Arsovski, A. Galstyan, J. M. Guseman, J. L. Nemhauser, Photomorphogenesis. The Arabidopsis Book 2012 (2012).

4. X. Huang, X. Ouyang, X. W. Deng, Beyond repression of photomorphogenesis: role switching of COP/DET/FUS in light signaling. Current Opinion in Plant Biology 21, 96–103 (2014).

5. X. W. Deng, T. Caspar, P. H. Quail, cop1: a regulatory locus involved in light-controlled development and gene expression in Arabidopsis. Genes Dev 5, 1172–1182 (1991).

6. D. Zhu et al., Biochemical characterization of Arabidopsis complexes containing CONSTITUTIVELY PHOTOMORPHOGENIC1 and SUPPRESSOR OF PHYA proteins in light control of plant development. Plant Cell 20, 2307–2323 (2008).

7. M. Holm, L. G. Ma, L. J. Qu, X. W. Deng, Two interacting bZIP proteins are direct targets of COP1-mediated control of light-dependent gene expression in Arabidopsis. Genes Dev 16, 1247–1259 (2002).

8. P. D. Duek, M. V. Elmer, V. R. van Oosten, C. Fankhauser, The Degradation of HFR1, a Putative bHLH Class Transcription Factor Involved in Light Signaling, Is Regulated by Phosphorylation and Requires COP1. Current Biology 14, 2296–2301 (2004).

9. I. C. Jang, J. Y. Yang, H. S. Seo, N. H. Chua, HFR1 is targeted by COP1 E3 ligase for post-translational proteolysis during phytochrome A signaling. Genes Dev 19, 593–602 (2005).

10. H. S. Seo et al., LAF1 ubiquitination by COP1 controls photomorphogenesis and is stimulated by SPA1. Nature 423, 995–999 (2003).

11. FAO (2024) Global status of salt-affected soils. (Rome, Italy).

12. P. Parihar, S. Singh, R. Singh, V. P. Singh, S. M. Prasad, Effect of salinity stress on plants and its tolerance strategies: a review. Environ Sci Pollut Res Int 22, 4056–4075 (2015).

13. E. van Zelm, Y. Zhang, C. Testerink, Salt Tolerance Mechanisms of Plants. Annu Rev Plant Biol 71, 403–433 (2020).

14. K. Atta et al., Impacts of salinity stress on crop plants: improving salt tolerance through genetic and molecular dissection. Frontiers in Plant Science Volume 14 - 2023 (2023).

15. C. Liu, X. Jiang, Z. Yuan, Plant Responses and Adaptations to Salt Stress: A Review. Horticulturae 10, 1221 (2024).

16. A. Ali, V. Petrov, D.-J. Yun, T. Gechev, Revisiting plant salt tolerance: novel components of the SOS pathway. Trends in Plant Science 28, 1060–1069 (2023).

17. L. Ma et al., Phytochromes enhance SOS2-mediated PIF1 and PIF3 phosphorylation and degradation to promote Arabidopsis salt tolerance. Plant Cell 35, 2997–3020 (2023).

18. X. Liu et al., FERONIA coordinates plant growth and salt tolerance via the phosphorylation of phyB. Nat Plants 9, 645–660 (2023).

19. P. Qi, W. Mo, R. Lin, The phytochrome B signaling regulates salt-mediated seedling growth in the dark. Plant and Cell Physiology 66, 766–780 (2025).

20. P. Žádníková et al., Role of PIN-mediated auxin efflux in apical hook development of Arabidopsis thaliana. Development 137, 607–617 (2010).

21. M. C. Cheng, P. K. Kathare, I. Paik, E. Huq, Phytochrome Signaling Networks. Annu Rev Plant Biol 72, 217–244 (2021).

22. H. Shi et al., Seedlings Transduce the Depth and Mechanical Pressure of Covering Soil Using COP1 and Ethylene to Regulate EBF1/EBF2 for Soil Emergence. Curr Biol 26, 139–149 (2016).

23. X. Shen, Y. Li, Y. Pan, S. Zhong, Activation of HLS1 by Mechanical Stress via Ethylene-Stabilized EIN3 Is Crucial for Seedling Soil Emergence. Frontiers in Plant Science 7 (2016).

24. K. Jonsson et al., Mechanochemical feedback mediates tissue bending required for seedling emergence. Curr Biol 31, 1154–1164 e1153 (2021).

25. J. Friml, J. Wiśniewska, E. Benková, K. Mendgen, K. Palme, Lateral relocation of auxin efflux regulator PIN3 mediates tropism in Arabidopsis. Nature 415, 806–809 (2002).

26. F. Vandenbussche et al., The auxin influx carriers AUX1 and LAX3 are involved in auxin-ethylene interactions during apical hook development in Arabidopsis thaliana seedlings. Development 137, 597–606 (2010).

27. L. Jo, K. Kajala, ggPlantmap: an open-source R package for the creation of informative and quantitative ggplot maps derived from plant images. J Exp Bot 75, 5366–5376 (2024).

28. F. Lin et al., B-BOX DOMAIN PROTEIN28 Negatively Regulates Photomorphogenesis by Repressing the Activity of Transcription Factor HY5 and Undergoes COP1-Mediated Degradation. Plant Cell 30, 2006–2019 (2018).

29. M. Ruan et al., Alternative oxidase 2 influences Arabidopsis seed germination under salt stress by modulating ABA signalling and ROS homeostasis. Environmental and Experimental Botany 217, 105568 (2024).

30. Y. Zou, Y. Zhang, C. Testerink, Root dynamic growth strategies in response to salinity. Plant Cell Environ 45, 695–704 (2022).

31. N. Gigli-Bisceglia, E. van Zelm, W. Huo, J. Lamers, C. Testerink, Arabidopsis root responses to salinity depend on pectin modification and cell wall sensing. Development 149 (2022).

32. H. J. Park et al., S-acylated and nucleus-localized SALT OVERLY SENSITIVE3/CALCINEURIN B-LIKE4 stabilizes GIGANTEA to regulate Arabidopsis flowering time under salt stress. Plant Cell 35, 298–317 (2023).

33. Y. Zhang, N. Li, L. Wang, Phytochrome interacting factor proteins regulate cytokinesis in Arabidopsis. Cell Rep 35, 109095 (2021).

34. D. Xu, COP1 and BBXs-HY5-mediated light signal transduction in plants. New Phytologist 228, 1748–1753 (2020).

35. A. Yadav, N. Ravindran, D. Singh, P. V. Rahul, S. Datta, Role of Arabidopsis BBX proteins in light signaling. Journal of Plant Biochemistry and Biotechnology 29, 623–635 (2020).

36. J. Cao et al., Multi-layered roles of BBX proteins in plant growth and development. Stress Biol 3, 1 (2023).

37. S. Li et al., Genome-wide identification of B-box zinc finger (BBX) gene family in Medicago sativa and their roles in abiotic stress responses. BMC Genomics 25, 110 (2024).

38. S. Nagaoka, T. Takano, Salt tolerance-related protein STO binds to a Myb transcription factor homologue and confers salt tolerance in Arabidopsis. Journal of Experimental Botany 54, 2231–2237 (2003).

39. T. S. Chiriotto, M. Saura-Sanchez, C. Barraza, J. F. Botto, BBX24 Increases Saline and Osmotic Tolerance through ABA Signaling in Arabidopsis Seeds. Plants (Basel) 12 (2023).

40. S. K. Meena et al., Antisense transcription from stress-responsive transcription factors fine-tunes the cold response in Arabidopsis. The Plant Cell 36, 3467–3482 (2024).

41. Y. Ma et al., Endoreplication mediates cell size control via mechanochemical signaling from cell wall. Science Advances 8, eabq2047 (2022).

42. H. Ren, M. Y. Park, A. K. Spartz, J. H. Wong, W. M. Gray, A subset of plasma membrane-localized PP2C.D phosphatases negatively regulate SAUR-mediated cell expansion in Arabidopsis. PLOS Genetics 14, e1007455 (2018).

43. J. Wang et al., SAUR17 and SAUR50 Differentially Regulate PP2C-D1 during Apical Hook Development and Cotyledon Opening in Arabidopsis. The Plant Cell 32, 3792–3811 (2020).

44. C. Costigliolo Rojas et al., Organ-specific COP1 control of BES1 stability adjusts plant growth patterns under shade or warmth. Dev Cell 57, 2009–2025 e2006 (2022).

45. S. Hayes et al., Warm temperature and mild water stress cooperatively promote root elongation. Curr Biol 10.1016/j.cub.2025.06.062 (2025).

46. X. Han et al., Time series single-cell transcriptional atlases reveal cell fate differentiation driven by light in Arabidopsis seedlings. Nat Plants 9, 2095–2109 (2023).

47. Z. Liu et al., Identification of the Regulators of Epidermis Development under Drought- and Salt-Stressed Conditions by Single-Cell RNA-Seq. Int J Mol Sci 23 (2022).

48. W. Wang, Q. Chen, J. R. Botella, S. Guo, Beyond Light: Insights Into the Role of Constitutively Photomorphogenic1 in Plant Hormonal Signaling. Front Plant Sci 10, 557 (2019).

49. Y. Liu, Y. Xie, D. Xu, X. W. Deng, J. Li, Inactivation of GH3.5 by COP1-mediated K63-linked ubiquitination promotes seedling hypocotyl elongation. Nat Commun 16, 3541 (2025).

50. M. Pacín, M. Semmoloni, M. Legris, S. A. Finlayson, J. J. Casal, Convergence of CONSTITUTIVE PHOTOMORPHOGENESIS 1 and PHYTOCHROME INTERACTING FACTOR signalling during shade avoidance. New Phytologist 211, 967–979 (2016).

51. S. N. Gangappa, S. V. Kumar, DET1 and HY5 Control PIF4-Mediated Thermosensory Elongation Growth through Distinct Mechanisms. Cell Rep 18, 344–351 (2017).

52. M. Sassi et al., COP1 mediates the coordination of root and shoot growth by light through modulation of PIN1- and PIN2-dependent auxin transport in Arabidopsis. Development 139, 3402–3412 (2012).

53. M. J. Iglesias et al., Shade-induced ROS/NO reinforce COP1-mediated diffuse cell growth. Proc Natl Acad Sci U S A 121, e2320187121 (2024).

54. T. W. McNellis et al., Genetic and molecular analysis of an allelic series of cop1 mutants suggests functional roles for the multiple protein domains. The Plant Cell 6, 487–500 (1994).

55. J. W. Reed, P. Nagpal, D. S. Poole, M. Furuya, J. Chory, Mutations in the gene for the red/far-red light receptor phytochrome B alter cell elongation and physiological responses throughout Arabidopsis development. The Plant Cell 5, 147–157 (1993).

56. J. W. Reed, A. Nagatani, T. D. Elich, M. Fagan, J. Chory, Phytochrome A and Phytochrome B Have Overlapping but Distinct Functions in Arabidopsis Development. Plant Physiol 104, 1139–1149 (1994).

57. B. C. Willige et al., D6PK AGCVIII Kinases Are Required for Auxin Transport and Phototropic Hypocotyl Bending in Arabidopsis The Plant Cell 25, 1674–1688 (2013).

58. J.J. Küpers et al., Local light signaling at the leaf tip drives remote differential petiole growth through auxin-gibberellin dynamics. Current Biology 33, 75-85.e75 (2023).

59. E. van Zelm et al., Natural variation in salt-induced root growth phases and their contribution to root architecture plasticity. Plant, Cell & Environment 46, 2174–2186 (2023).

60. J. Schindelin et al., Fiji: an open-source platform for biological-image analysis. Nature Methods 9, 676–682 (2012).

61. Q. Zhu, P. Žádníková, D. Smet, D. Van Der Straeten, E. Benková, “Real-Time Analysis of the Apical Hook Development” in Plant Hormones: Methods and Protocols, J. Kleine-Vehn, M. Sauer, Eds. (Springer New York, New York, NY, 2017), 10.1007/978-1-4939-6469-7_1, pp. 1-8.

62. J. Stuttmann et al., Highly efficient multiplex editing: one-shot generation of 8× Nicotiana benthamiana and 12× Arabidopsis mutants. Plant J 106, 8–22 (2021).

63. K. Labun et al., CHOPCHOP v3: expanding the CRISPR web toolbox beyond genome editing. Nucleic Acids Res 47, W171–w174 (2019).

64. D. Kurihara, Y. Mizuta, Y. Sato, T. Higashiyama, ClearSee: a rapid optical clearing reagent for whole-plant fluorescence imaging. Development 142, 4168–4179 (2015).

65. J. J. Kupers et al., Local light signaling at the leaf tip drives remote differential petiole growth through auxin-gibberellin dynamics. Curr Biol 33, 75–85 e75 (2023).

66. D. Swarbreck et al., The Arabidopsis Information Resource (TAIR): gene structure and function annotation. Nucleic Acids Research 36, D1009–D1014 (2008).

67. P. A. Ewels et al., The nf-core framework for community-curated bioinformatics pipelines. Nature Biotechnology 38, 276–278 (2020).

68. M. I. Love, W. Huber, S. Anders, Moderated estimation of fold change and dispersion for RNA-seq data with DESeq2. Genome Biol 15, 550 (2014).

69. A. Zhu, J. G. Ibrahim, M. I. Love, Heavy-tailed prior distributions for sequence count data: removing the noise and preserving large differences. Bioinformatics 35, 2084–2092 (2018).

70. T. Wu et al., clusterProfiler 4.0: A universal enrichment tool for interpreting omics data. The Innovation 2, 100141 (2021).

