## Supplemental Information for "Salt stress disrupts local auxin and COP1 gradients in Arabidopsis apical hooks"

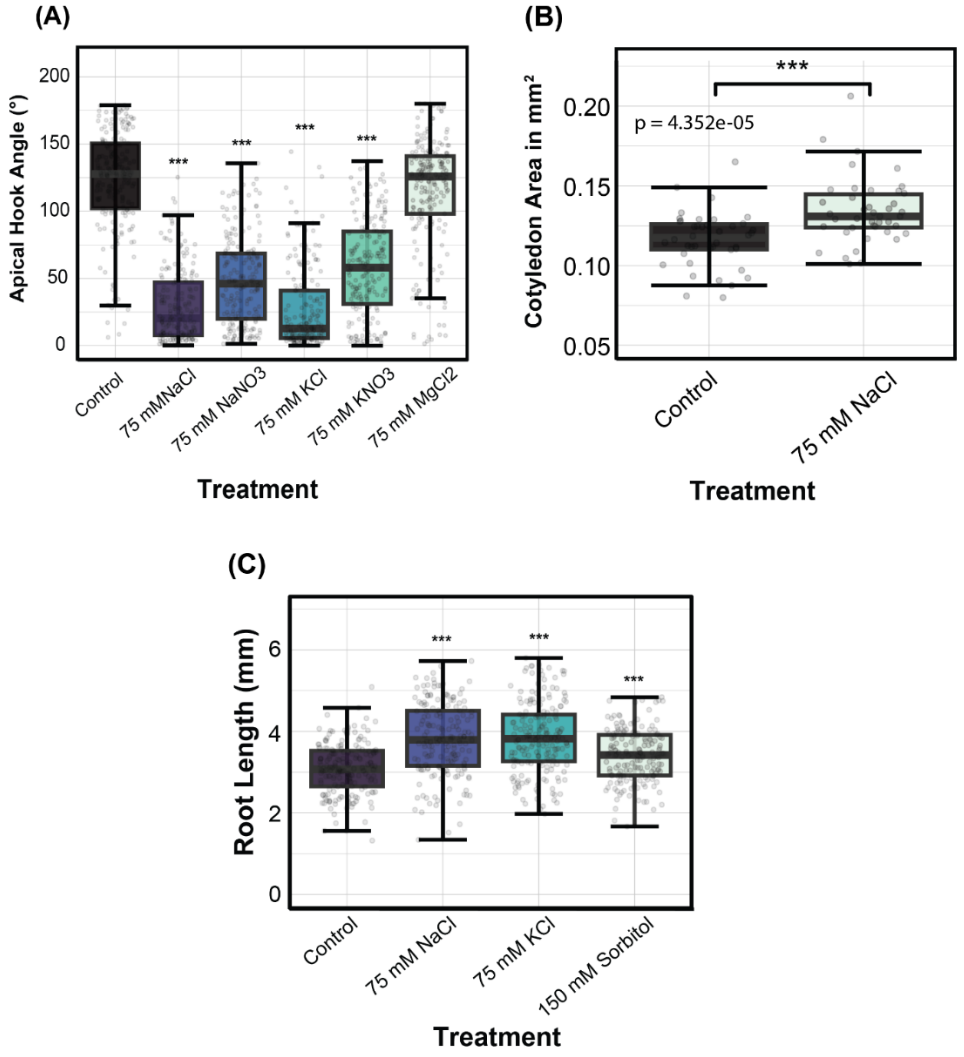


***Supplementary Figure 1. Salinity induces a mild photomorphogenic phenotype in etiolated seedlings – continued.*** (A) Boxplot representing apical hook angles of three-day old dark grown seedlings treated with 75 mM of NaCl, $NaNO_3$, KCl, $KNO_3$ and $MgCl_2$. (B) Boxplot representing cotyledon size for three-day old dark grown seedlings treated with 75 mM NaCl and control. (C) Boxplot representing root lengths of three-day old dark grown seedlings treated with 75 mM of NaCl, KCl and 150 Mm sorbitol. All boxplots represent the median ± 25% and 50%. Asterisks for all panels refer to significant differences compared to control (* $P \leq 0.05$, ** $P \leq 0.05$, *** $P \leq 0.001$) calculated via Kruskal Wallis followed by posthoc Dunn's test (A), or using independent samples t-test (B) or Welch's ANOVA followed by post-hoc Games-Howell test (C). dAll panels refer to phenotype data where seeds germinated on 0.5 MS plates, by exposure to ~125 µmol/m2/s white light for one hour followed by a further 23 hours in dark before being transferred to respective treatment media.

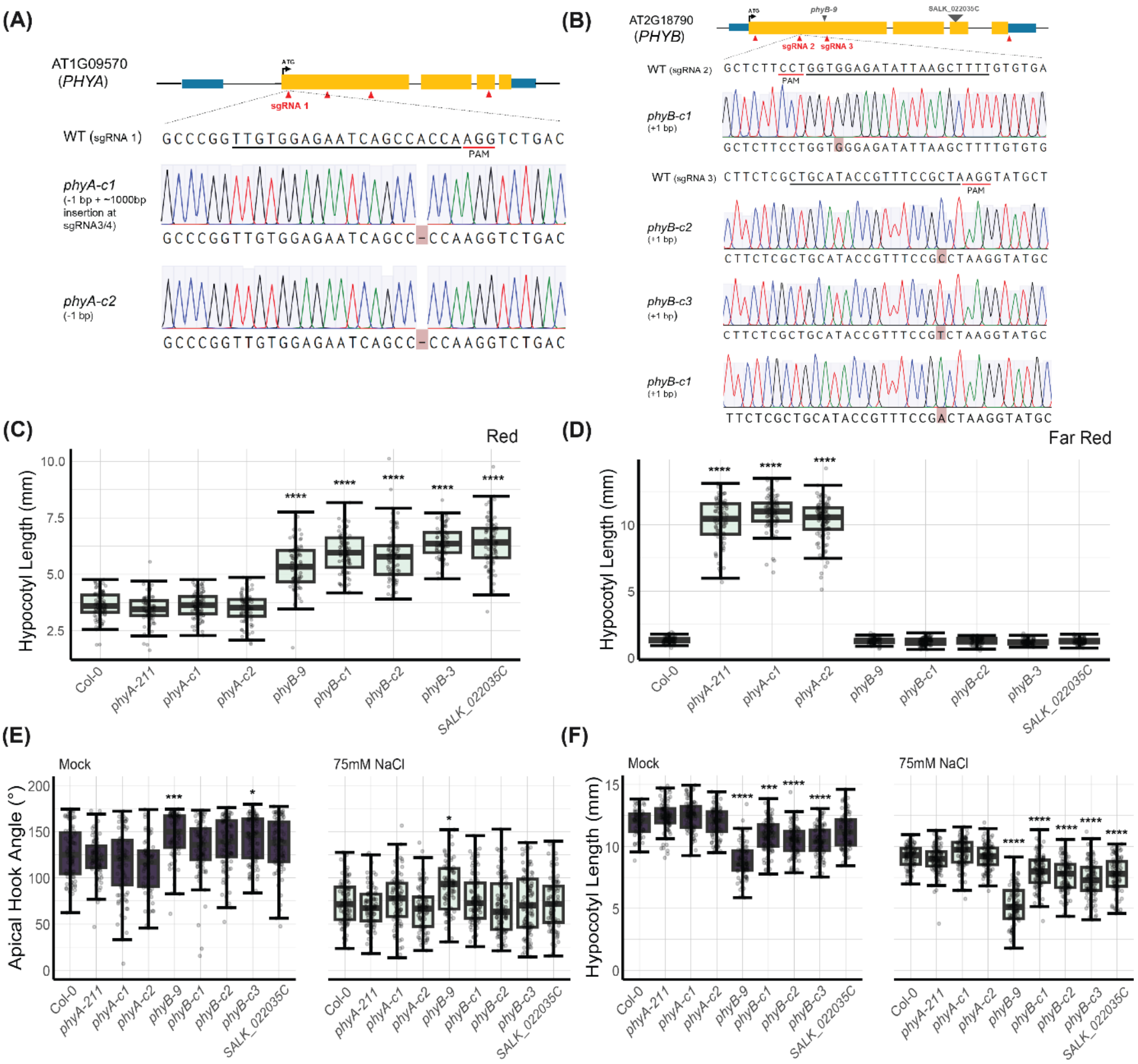


***Supplementary Figure 2.*** Mutations in three independent *phyA* (A) and *phyB* (B) alleles generated using CRISPR/Cas9 method compared to EMS (*phyb-9*), γ-irradiation & (*phyA-211*) and SALK T-DNA insertion *phyB* mutant (SALK_022035C). Red arrows indicate sgRNA target locations. White box indicates deletion, dashes refer to single nucleotide deletions, and highted letters refer to single nucleotide insertions. Boxplots representing hypocotyl lengths of different *phyA* and *phyB* mutants grown under red light for three days (C) or darkness for one day, followed by far red for two days. Apical hook angle (E) and hypocotyl length (F) of three day old dark grown seedlings treated with 75 mM NaCl. All boxplots represent the median ± 25% and 50%. Asterisks for panels (C - F) refer to significant differences compared to control (* $P \leq 0.05$, ** $P \leq 0.05$, *** $P \leq 0.001$) calculated via Kruskal Wallis followed by posthoc Dunn's test Panels C-F refer to phenotype data where germination was induced by exposure to ~125 $\mu mol/m^2/s$ white light for three hours, before being transferred to 10 $\mu mol/m^2/s$ red light for three days (C), or dark for a further 23 hours (D-F). Dark germinated seeds were then placed in 10 $\mu mol/m^2/s$ far red light (D) or transferred to respective treatment media (75 mM NaCl / Control) for a further two days.

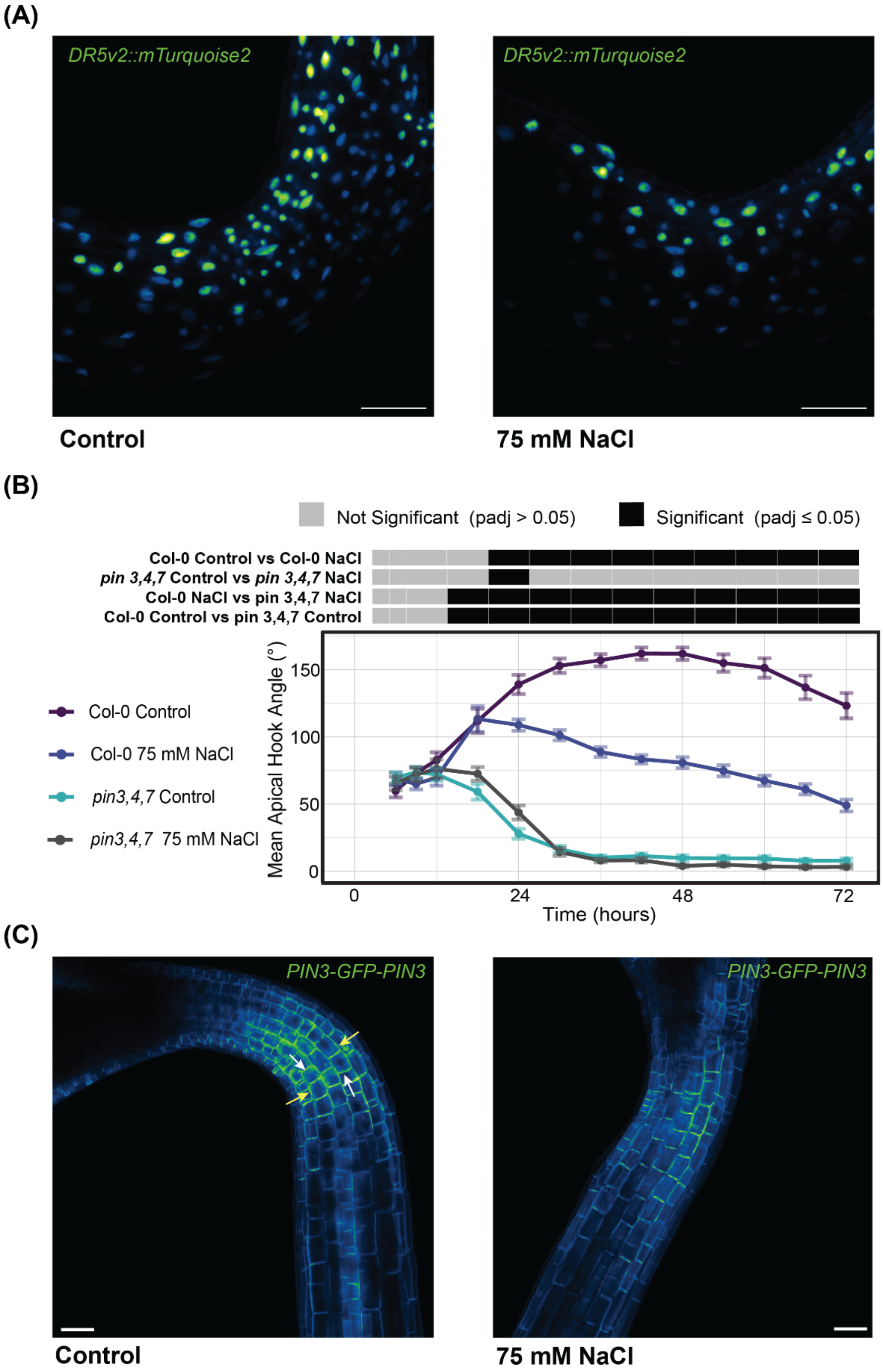


***Supplementary Figure 3.*** (A) Representative confocal images of *DR5v2::mTurqoise2* in the apical hooks of three day old etiolated seedlings treated with 75 mM NaCl or grown on control media. (B) Time-lapse data for apical hook angle following transfer (24h), where points represent mean angle, error bars represent ± SE, n ≥30. Significance calculated using pairwise t-test with Benjamini-Hochberg (BH) correction for multiple comparisons. (C) Representative confocal images of *PIN3-*

*GFP-PIN3* in two day old etiolated seedlings treated with 75 mM NaCl or grown on control media. White arrows point to anticlinal cell walls and yellow arrows point to periclinal cell walls used for quantifications. All panels refer to seedlings where seeds germinated on 0.5 MS plates, by exposure to ~125 μmol/m2/s white light for one hour followed by a further 23 hours in dark before being transferred to respective treatment media. Scale bars = 50 μm.

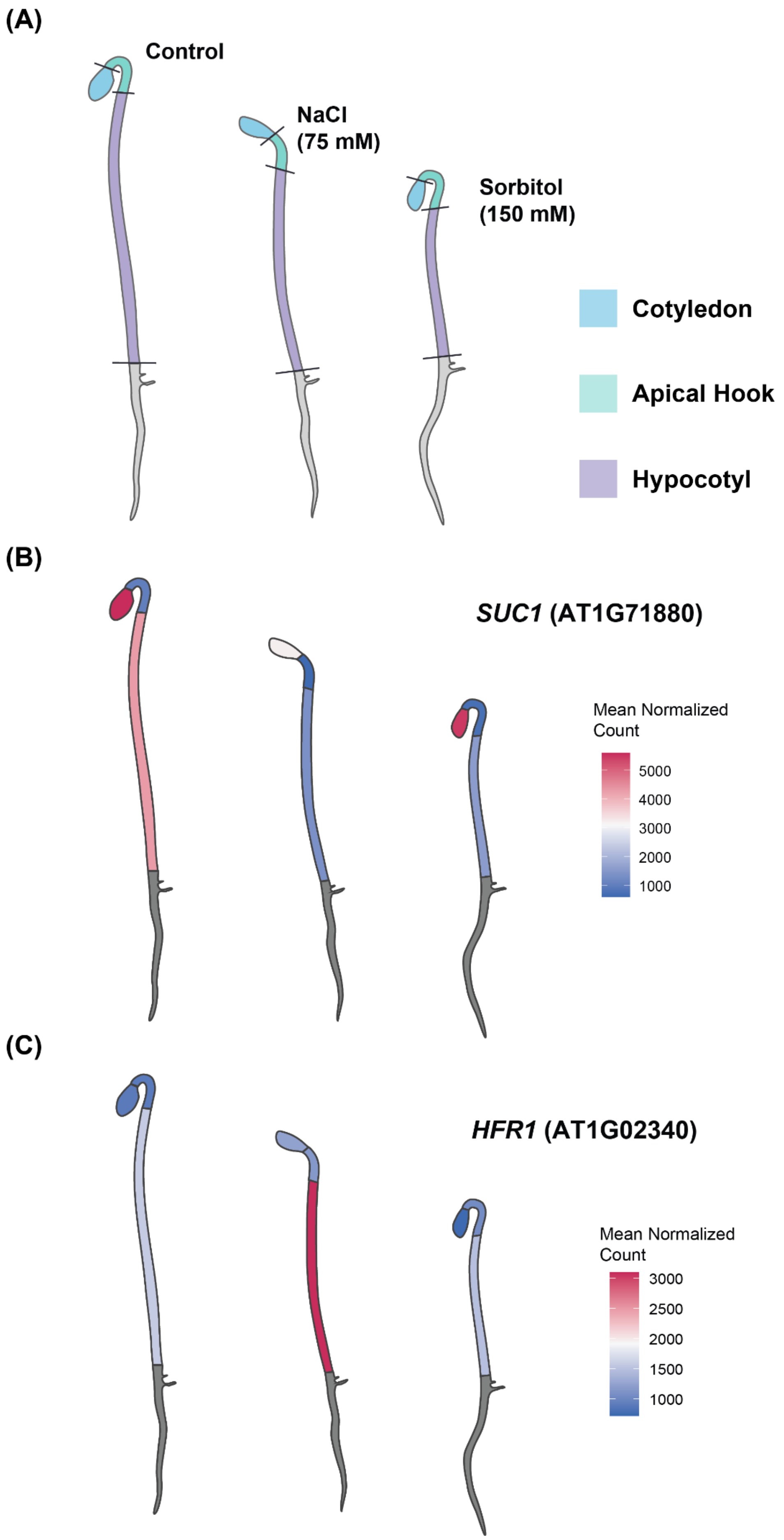
(A)
Control
NaCl
(75 mM)
Sorbitol
(150 mM)
Cotyledon
Apical Hook
Hypocotyl
(B)
SUC1 (AT1G71880)
Mean Normalized
Count
5000
4000
3000
2000
1000
(C)
HFR1 (AT1G02340)
Mean Normalized
Count
3000
2500
2000
1500
1000

***Supplementary Figure 4.*** (A) Representative seedling dissection diagram for RNA-sequencing of cotyledons, apical hooks, and hypocotyls. (B & C) *GGPlantmap* expression map showing relative transcript abundance (DESeq2 normalized reads) of *SUC1* and *HFR1*.

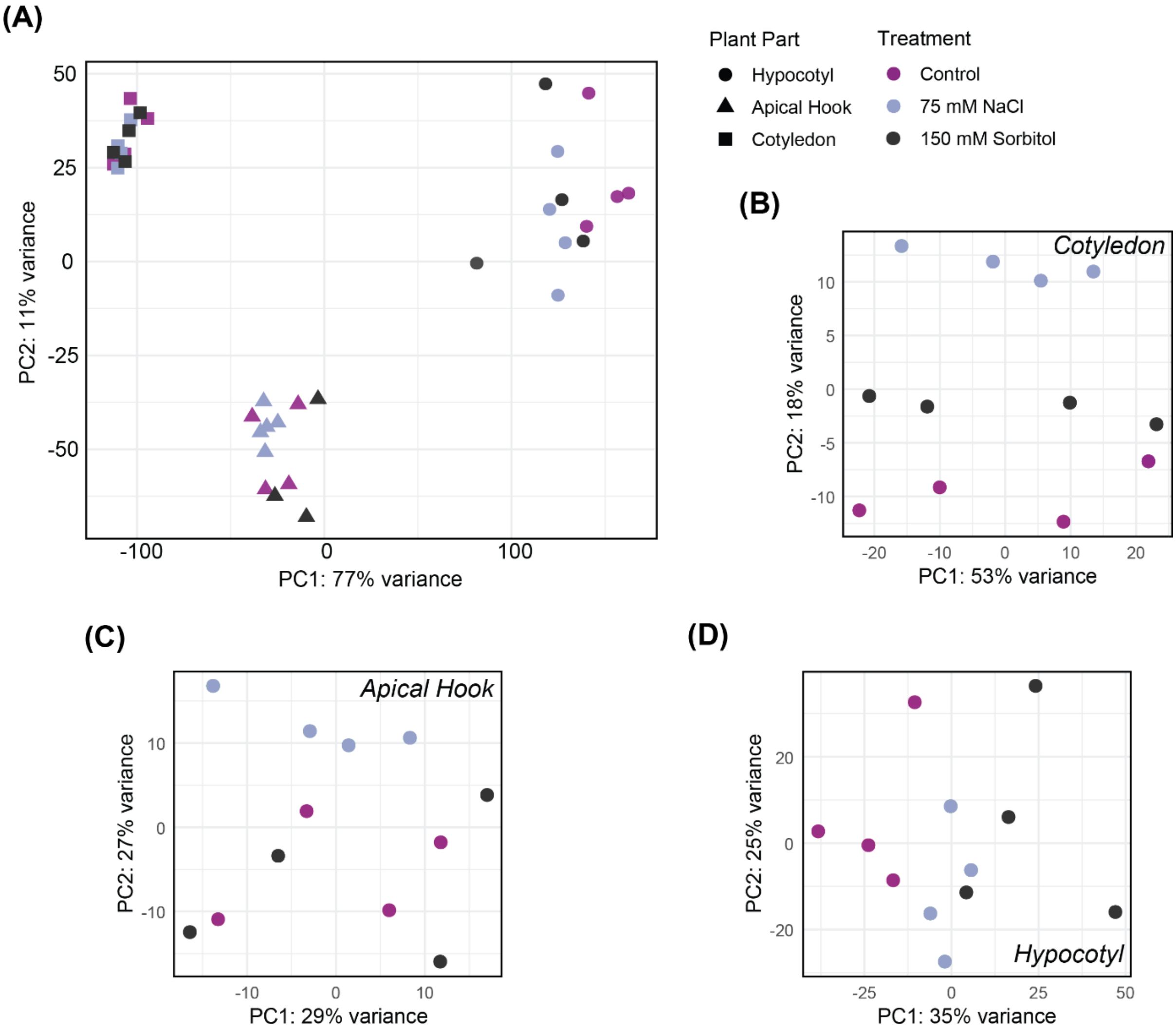


***Supplementary Figure 5.*** Principal component analysis (PCA) of relative log expression (RLD)–transformed counts for the top 10,000 most variable genes from RNA-seq data. (A) PCA of all samples from cotyledons, apical hooks, and hypocotyls under control, NaCl, and sorbitol treatments. Points are coloured by treatment and shaped by plant part. (B–D) PCA plots for each organ separately: (B) cotyledons, (C) apical hooks, and (D) hypocotyls, showing treatment-specific transcriptomic variation within each organ. Axes represent principal components 1 and 2, with the percentage of variance explained indicated. Each point represents one biological replicate.

***Supplementary Table 4.*** Oligos used to assemble shuttle and transformation vectors targeting *PHYA* and *PHYB*.

| Target gene | Shuttle vector | Forward oligo | Reverse oligo |
|---|---|---|---|
| *PHYA* [AT1G09570] | M1 pDGE332 | YHT046: attgAGGCTGCAATCTTTACCCAG | YHT047: aaacCTGGGTAAAGATTGCAGCCT |
| | M2 pDGE333 | YHT048: attgTTGTGGAGAATCAGCCACCA | YHT049: aaacTGGTGGCTGATTCTCCACAA |
| | M3 pDGE335 | YHT050: attgGTGGAACTCGATAACCAGA | YHT051: aaacTCTGGTTATCGAGTTCCAC |
| | M4E pDGE337 | YHT052: attgTAACTGTTTCAGCTTCCCTG | YHT053: aaacCAGGGAAGCTGAAACAGTTA |
| *PHYB* [AT2G18790] | M1 pDGE332 | YHT054: attgTTAGGAGTGTGACTTGACGA | YHT055: aaacTCGTCAAGTCACACTCCTAA |
| | M2 pDGE333 | YHT056: attgAAAAGCTTAATATCTCCACC | YHT057: aaacGGTGGAGATATTAAGCTTTT |
| | M3 pDGE335 | YHT058: attgCTGCATACCGTTTCCGCTA | YHT059: aaacTAGCGGAAACGGTATGCAG |
| | M4E pDGE337 | YHT060: attgAATACATCCGAGAATCAGAA | YHT061: aaacTTCTGATTCTCGGATGTATT |

***Supplementary Table 5.*** Genotyping primers used to screen for mutations and absence of Cas9.

| Target gene | Primers |
|---|---|
| *PHYA* [AT1G09570] | YHT084: CTAGGCCGACTCAGTCCTCT |
| | YHT085: TTGTTTGCTGCAGCGAGTTC |
| *PHYB* [AT2G18790] | YHT088: GGCACACATTTTGCTTCGTCT |
| | YHT089: ACAATGGTCGCTTTCGAGGT |
| Cas9 | JS184: CGAAGTTCCAAGGCGTGATA |
| | JS185: TAGGTGCAAGTCAGGAGGAA |
